# Single cell profiling of T and B cell repertoires following SARS-CoV-2 mRNA vaccine

**DOI:** 10.1101/2021.07.14.452381

**Authors:** Suhas Sureshchandra, Sloan A. Lewis, Brianna Doratt, Allen Jankeel, Izabela Ibraim, Ilhem Messaoudi

**Affiliations:** Department of Molecular Biology and Biochemistry, University of California, Irvine CA 92697; Institute for Immunology, University of California, Irvine CA 92697; Center for Virus Research, University of California, Irvine CA 92697

**Keywords:** SARS-CoV-2, mRNA vaccine, BCR, TCR, COVID-19

## Abstract

mRNA based vaccines for SARS-CoV-2 have shown exceptional clinical efficacy providing robust protection against severe disease. However, our understanding of transcriptional and repertoire changes following full vaccination remains incomplete. We used single-cell RNA sequencing and functional assays to compare humoral and cellular responses to two doses of mRNA vaccine with responses observed in convalescent individuals with asymptomatic disease. Our analyses revealed enrichment of spike-specific B cells, activated CD4 T cells, and robust antigen-specific polyfunctional CD4 T cell responses in all vaccinees. On the other hand, CD8 T cell responses were both weak and variable. Interestingly, clonally expanded CD8 T cells were observed in every vaccinee, as observed following natural infection. TCR gene usage, however, was variable, reflecting the diversity of repertoires and MHC polymorphism in the human population. Natural infection induced expansion of larger CD8 T cell clones occupied distinct clusters, likely due to the recognition of a broader set of viral epitopes presented by the virus not seen in the mRNA vaccine. Our study highlights a coordinated adaptive immune response where early CD4 T cell responses facilitate the development of the B cell response and substantial expansion of effector CD8 T cells, together capable of contributing to future recall responses.

## INTRODUCTION

The COVID-19 pandemic has spurred the rapid development of vaccines targeting SARS-CoV-2 that have garnered emergency use authorizations from the FDA and are being widely distributed (1, 2). The Pfizer (BNT162b2) and Moderna (mRNA-1273) mRNA-based vaccines were the first to be approved and have proven to be safe and efficacious (94% effective) in adults and children over 12 years of age (3, 4). However, the mechanisms by which these vaccines elicit long lasting cellular immune responses to SARS-CoV-2 and their mechanisms of action remain poorly understood. In this study, we aimed to address two questions: What are the functional and transcriptomic responses of memory T and B cells to mRNA-based COVID-19 vaccination? And how does the vaccine response differ from that of an asymptomatic SARS-CoV-2 infection?

Recent studies have shown that development of robust neutralizing antibody and memory B cell responses require both mRNA vaccine doses in SARS-CoV-2 naive individuals while comparable humoral responses are generated with just one dose in convalescent subjects (5). However, the ratio of binding to neutralizing antibodies after vaccination was greater than that after infection (6). Additionally, most vaccinees had Th1-skewed T cell responses where early Tfh and Th1 CD4 responses correlate with effective neutralizing antibody responses after the first dose and CD8 effector responses after the second dose (7). Furthermore, expanded T cell clones detected following vaccination were predominantly memory cells whereas those detected during infection were to effector cells with acute infection (8). Collectively, these observations suggest distinct T and B cell responses following vaccination in comparison to natural infection. Additional studies that integrate functional, transcriptional, and repertoire analysis of the memory immune cell response to COVID-19 mRNA vaccination are needed (7).

In this study, we assayed humoral and cellular responses to two doses of mRNA vaccine (14 days post dose 2) in four individuals and compared parallel changes in their immune repertoire with changes observed in three convalescent individuals who experienced asymptomatic/mild COVID-19 (~30 days after positive COVID test). Single dose of vaccine induced neutralizing titers comparable to levels seen following asymptomatic/mild SARS-CoV-2 infection. However, neutralizing titers in vaccinees increased several fold following second vaccine dose exceeding those detected following asymptomatic/mild SARS-CoV-2 infection. Antigen-specific B cells were detected after the second dose. Single cell analysis revealed an expansion of activated CD4 T cells but not CD8 T cells post vaccination. Robust antigen specific polyfunctional CD4 T cell responses were observed in all vaccinated individuals. Although CD8 T cell responses were weak and highly variable, effector memory CD8 T cell clones were expanded in every individual following vaccination. TCR gene usage was variable reflecting the diversity of T cell repertoires and MHC polymorphism in human population. Natural infection induced expansion of larger CD8 T cell clones, including distinct clusters likely due to the recognition of a broader set of epitopes presented by the virus not seen in the mRNA vaccine.

## RESULTS

### Humoral responses to SARS-CoV-2 mRNA vaccination

To comprehensively assess the cellular and humoral immune response to COVID-19 vaccination, we collected blood from SARS-CoV-2 naive volunteers prior to mRNA vaccination (baseline), and 2 weeks following prime-boost vaccination (post-vaccination dose 2; n=4) (**Figure 1A**). These responses were compared to those generated by individuals who experienced asymptomatic SARS-CoV-2 infection using longitudinal samples collected before (baseline) and ~30 days after exposure (convalescent, n=3). The demographics and vaccine information are provided in **Supplementary Table 1**. Both infection and vaccination induced binding **(Supp Figure 1A)** and neutralizing antibodies **(Supp Figure 1B)**, as early as 2 weeks following the first dose of vaccination that increased several folds following booster vaccination **(Figure 1B)** and reached slightly higher levels than those achieved following asymptomatic/mild infection (p=0.09). Given that full protection against the virus is achieved two weeks after the booster, we chose this time point for the additional downstream analyses.

**Figure 1:**
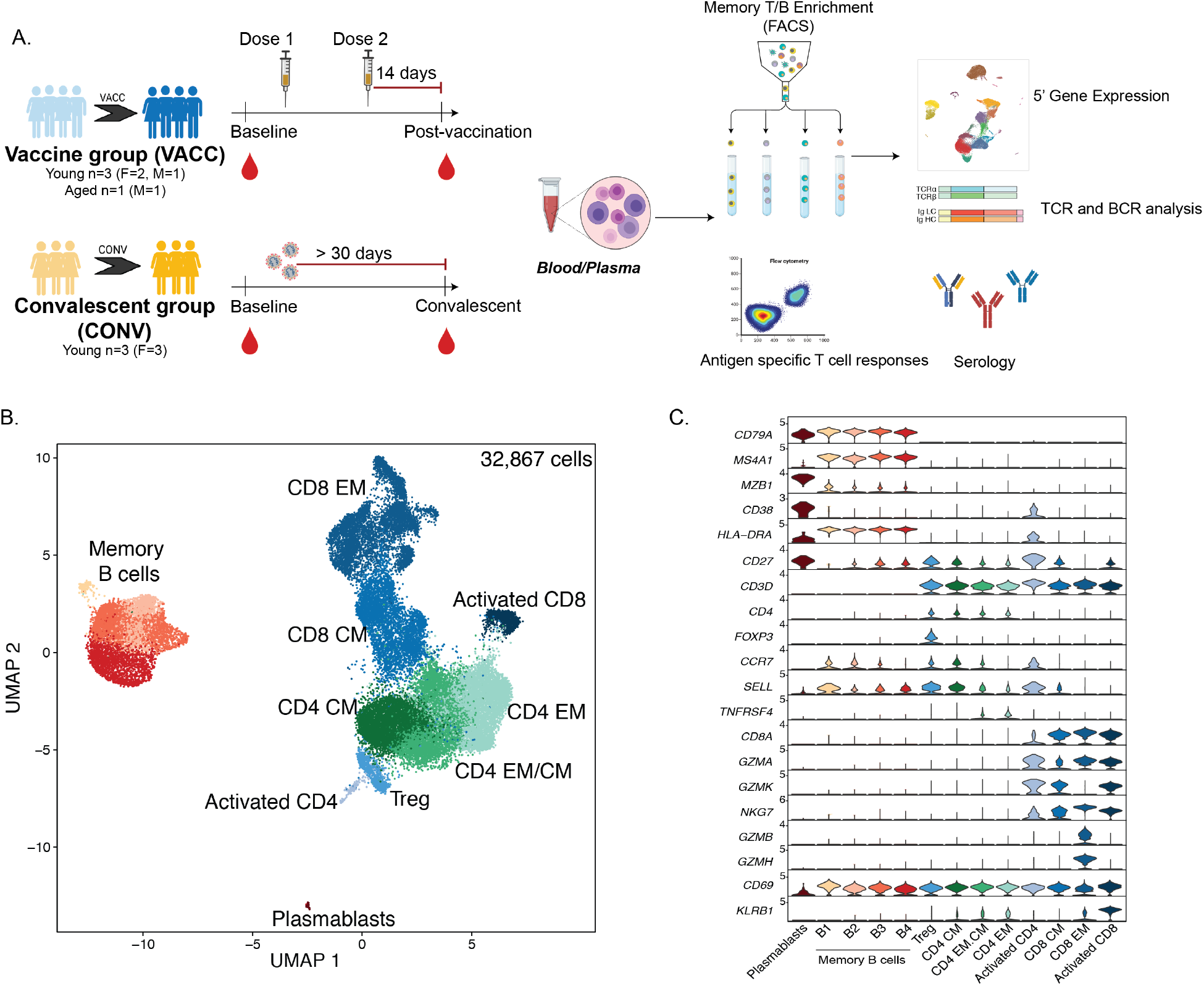
Immunological changes with SARS-CoV-2 mRNA vaccination. (A) Experimental design for the study. Blood was collected from SARS-CoV-2 naive subjects before vaccination, 2 weeks post dose 1, and 2 weeks post prime boost vaccination (VACC group) or in SARS-CoV-2 exposed but asymptomatic individuals (CONV group) before and after convalescence. Immune phenotypes of PBMC and antigen specific T and B cell responses were measured using multi-color flow cytometry. Longitudinal serological responses to the vaccine were measured using ELISA and neutralization assays. Memory T and B cells from a subset of PBMC samples (n=4/group for vaccine volunteers, n=3/group for convalescent health care workers, matched) were profiled using scRNA-Seq at the baseline (pre-vaccination or and post-vaccination time points. (B) UMAP projection of 32,867 memory T and B cells with major subsets annotated. (C) Violin plots of key gene markers used for cluster annotations. Normalized transcript counts are shown on the Y-axis. Figure 2: B cell adaptations following SARS-CoV-2 mRNA vaccination and infection

To specifically assess distinct memory responses, we sorted memory T and B cells, and circulating plasmablasts from PBMC before and 2 weeks after booster vaccination or ~30 days after SARS-CoV-2 infection **(Supp Figure 1C)** and performed 5’ scRNA-Seq combined with parallel repertoire analysis **(Figure 1A)**. Dimension reduction of 32,867 cells from 4 vaccinated and 3 convalescent individuals **(Supp Figure 1D)** by Uniform Manifold Approximation and Projection (UMAP) separated clusters of cells that were identified as regulatory T cells (*FOXP3*), effector memory (EM), central memory (CM), and activated CD4 and CD8 T cells **(Figures 1B and 1C)**. CM and EM subsets were distinguished based on relative expression of *CCR7, TNFRSF4, GZMH/B, NKG7*, and *SELL* (encoding CD62L), whereas activated CD4 T cells expressed high levels of *CD38*, *HLA-DR* and activated CD8 T cells expressed high levels of *CD69* and *KLRB1* **(Figure 1C)**. We also identified 4 subsets of memory B cells based on relative expression of *CD27*, *SELL*, and *CCR7*. A small cluster of plasmablasts was identified based on *MZB1* and *CD38* expression **(Figures 1B and 1C)**.

### B cell responses to vaccination and infection

We next examined the response to vaccination within the B cell compartment. Both vaccination and asymptomatic infection resulted in the reduction of naïve and expansion of memory B cells, with these changes being more prominent with natural infection **(Supp Figure 2A)**. Importantly, antigen (spike) specific B cells were detected in circulation two weeks after prime-boost vaccination **(Supp Figure 2B and Figure 2A)**. Examination of memory B cells using single cell RNA sequencing revealed four major clusters **(Figure 2B)** exhibiting distinct patterns of immunoglobulin genes **(Supp Figure 2C)**: 1) a less mature cluster B1 expressing high *IGHD* and *IGHM*; 2) cluster B2 expressing lower *IGHM* but higher *IGHA1*; 3) cluster B3 sharing features with B2 but also expressing *IGHG1* and *IGHG2*; and cluster B4 expressing highest levels of *IGHG2* **(Supp Figure 2C)**. Comparison of cluster proportions revealed a reduction in immature B1 cluster and enrichment of mature clusters B3 and B4 with both vaccination and infection **(Figure 2C)**. Modest increases in plasmablast frequencies were observed following vaccination **(Figure 2C and Supp Figure 2D).** We then compared gene expression profiles between memory B cells collected two weeks post vaccination and in convalescent patients ~30 days post exposure. Genes upregulated (Log_2_ Fold change ≥ 0.4 and FDR ≤ 0.05) with vaccination relative to infection (n=44) play a role in leukocyte activation while those upregulated with infection (n=96) enriched to additional GO terms including “cytokine production” revealed enrichment of leukocyte activation with both vaccination and convalescence, but over-representation of cytokine signaling, apoptosis, cell adhesion, and type-I IFN signaling exclusively with convalescence **(Supp Figure 2E)**.

**Figure 2:**
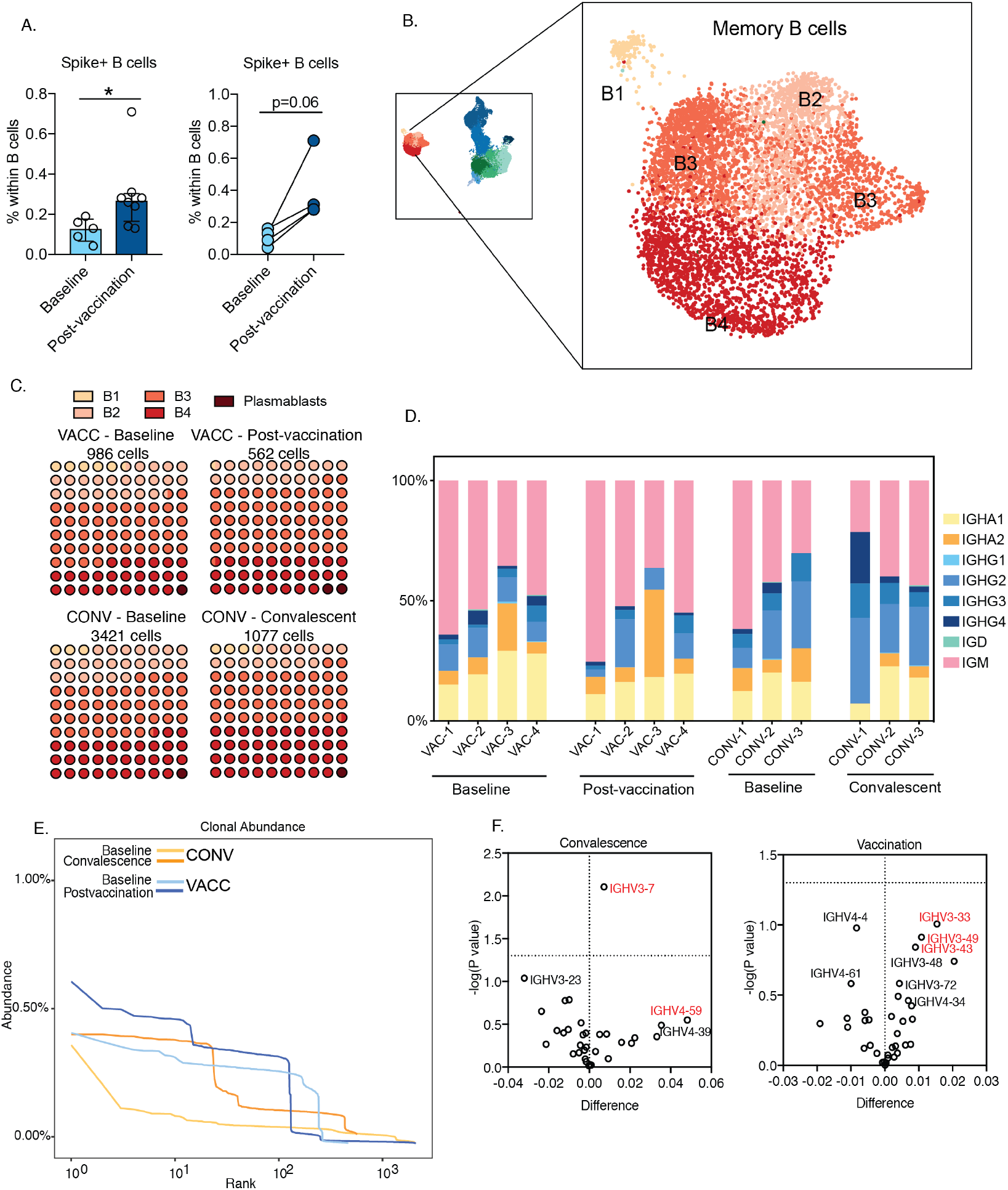
B cell adaptations following SARS-CoV-2 mRNA vaccination and infection. (A) Dot plots representing expansion of Spike+ cells within total CD20+ B cells in PBMC before and after vaccination (aggregate differences on the left and matched differences on the right). PBMC were incubated with biotinylated Spike protein and fluorochrome conjugated streptavidin, surface stained, washed and analyzed using flow cytometry. Group differences were tested using unpaired t-test with Welch’s correction (left panel) or paired t-test (right panel). (B) Magnified view of B cell subsets identified using single cell RNA sequencing. Data includes samples from all four groups. (C) Waffle plot representation of B cell cluster quantification with infection and vaccination. (D) Isotype distribution of productive B cell clones in vaccinated (n=4) and convalescent (n=3) individuals. Isotypes were determined based on the constant region of the clone. (E) Aggregate clonal abundance following vaccination and infection. (F) Volcano plots depicting heavy chain gene usage biases following convalescence (left panel, relative to pre-infection baseline) or vaccination (bottom panel, relative to pre vaccination baseline). X axis represents the change in gene usage and Y axis represents p-value (−log 10). Two-way comparisons were tested using ratio-paired test for matched comparisons, [p-values: *− p<0.05]

B cell repertoire analysis resolved memory B cells into distinct isotypes **(Figure 2D)**. Expansion of IgG+ cells was evident in a subset of individuals after vaccination and infection. Vaccination resulted in a significant reduction in IgA1+ memory B cells (p=0.018) **(Figure 2D)**. Clonal analysis of B cells revealed expansion of small size clones (10-100) with both vaccination and infection, albeit to a lesser magnitude with vaccination **(Figure 2E)**. Infection only was associated with expansion of larger B cell clones (>100) **(Figure 2E)**. Finally, gene usage analysis revealed preferential usage of heavy chain gene family IGHV3 - IGHV3-7, IGHV4-59 with convalescence and IGHV3-33, IGHV3-43, IGHV3-49 with vaccination **(Figure 2F)**.

### T cell adaptations with vaccination and convalescence

Next, we examined the effects of COVID-19 vaccination or natural infection on the distribution of memory T cell subsets. CD4 TCM subset expanded with vaccination (p=0.08) but not in convalescent subjects **(Supp Figure 3A)**. No major changes in other CD4 subsets (naïve, TEM) or CD8 subsets were detected with flow cytometry **(Supp Figures 3A and 3B)**. Single cell analyses revealed an expansion of activated CD4+ T cells **(Figure 3A)** with vaccination. This cluster of CD4 T cells expressed relatively higher levels of *CD38* and *HLA-DRA* and cytotoxic molecules *GZMK* and *PRF1* **(Figure 3B)**. On the other hand, frequencies of activated CD8 T cells (expressing *CD69* and *KLRG1*) were comparable across groups **(Figures 3A and 3B)**. The expansion of activated HLA-DR+CD38+ CD4 T cells following vaccination was confirmed using flow cytometry **(Figures 3C and 3D)**. Activated CD8 T cells on the other hand, exhibited increased RNA levels of activation markers *CD69* and *TNFAIP3* with both infection and vaccination **(Figure 3E)**, however, factors associated with memory development (*FOS*, *KLF6*) were up-regulated with vaccination and not convalescence **(Figure 3E)**. Within the CD8 EM subset, both vaccination and infection up-regulated *BCL3* **(Figure 3F)**, which is essential for maximum IFNγ secretion following secondary antigen stimulation. However, tissue homing factor *SELPG* (encoding P-selectin) and cytotoxic gene *GNLY* (encoding Granulysin) were induced only with vaccination **(Figure 3F)**.

**Figure 3:**
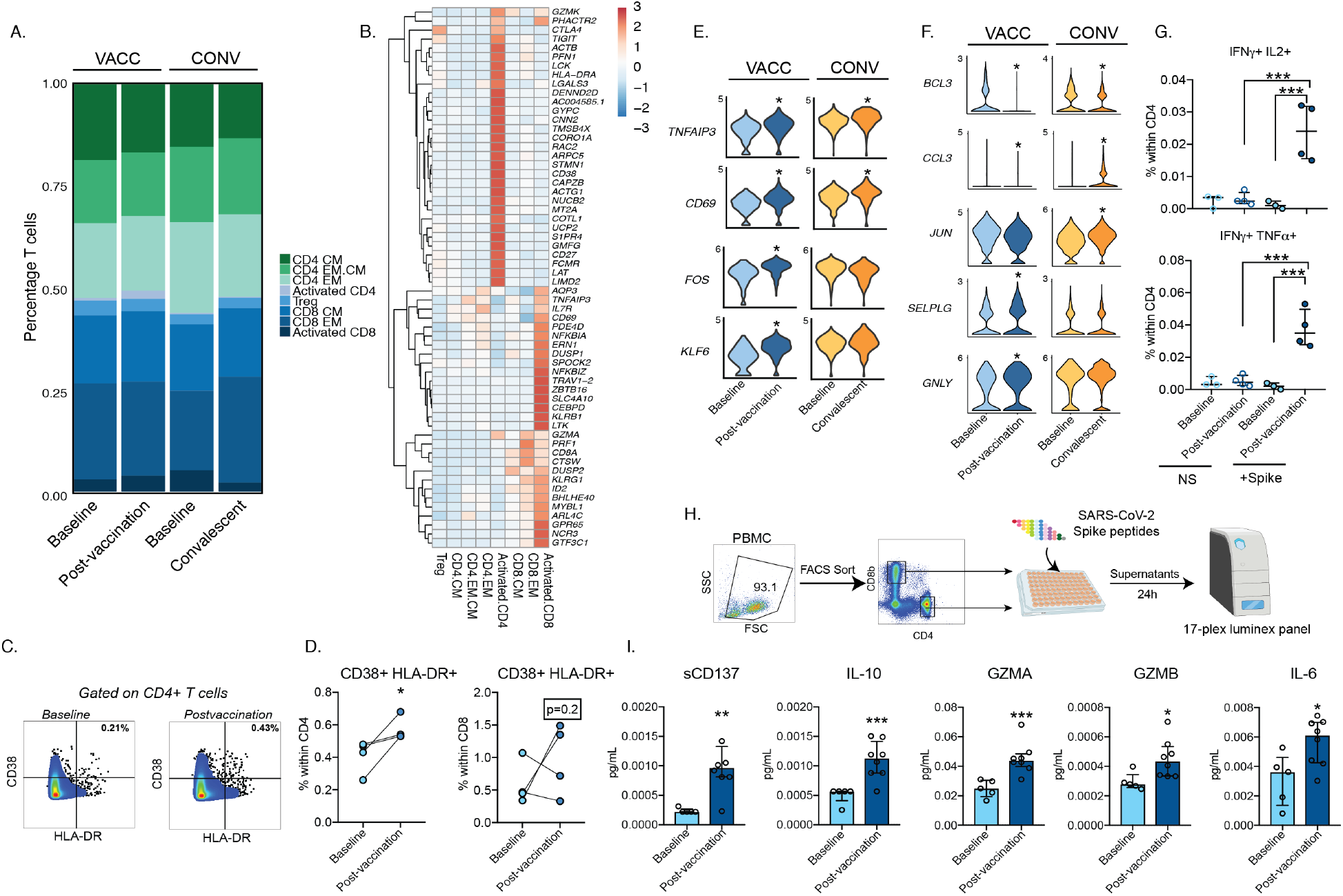
T cell adaptations with SARS-CoV-2 mRNA vaccination. (A) Stacked bar graph comparing the distribution of memory CD4 and CD8 T cells across each group reported as percentage total cells. (B) Clustered heatmap comparing aggregate top markers from each of the memory T cell clusters. Colors represent normalized transcript levels ranging from low (in blue) to high (in red). (C) Gating strategy for identification of activated CD4 and CD8 T cells before and after vaccination. (D) Frequencies of CD38+HLA-DR+ CD4 and CD8 T cells following vaccination. (E-F) Violin plots comparing key genes differentially expressed in (E) activated CD8 and (F) CD8 EM subsets either with convalescence and/or vaccination. (G) Polyfunctional CD4 T cell responses following overnight SARS-CoV-2 spike peptide stimulation measured using intracellular cytokine staining and flow cytometry. (FI) Experimental design for measuring T cell effector responses following vaccination. Sorted CD4 and CD8 T cells were stimulated with SARS-CoV-2 overlapping spike peptides for 24 hours. Supernatants were collected and analyzed using Luminex. (I) Secreted levels of soluble co-stimulatory molecule (sCD137), cytokines (IL-10, IL-6), and effector molecules (Granzyme A and Granzyme B) in CD4 T cells following mRNA vaccination. Two-way comparisons were tested using either ratio-paired test for matched comparisons or unpaired t-test with Welch’s correction for group comparisons. Four way comparisons were tested using one-way ANOVA followed by Flolm Sidak’s multiple hypothesis correction [p-values: *− p<0.05; **−p<0.01; ***−p<0.001; ****−p<0.0001]

### Robust antigen specific CD4 T cell effector responses with vaccination

We next interrogated antigen specific CD4 and CD8 T cell responses with vaccination. Total PBMC were stimulated with an overlapping peptides library covering the entire sequence of the spike protein for 24 hours, surface stained, fixed, and analyzed for cytokine production using flow cytometry **(Supp Figure 3C)**. Spike specific polyfunctional IFNγ+IL-2+ and IFNγ+TNFα+ CD4, but not CD8, T cells were evident 2 weeks post prime-boost vaccination **(Figure 3G and Supp Figure 3D)**. To gain a more comprehensive understanding of the effector T cell response induced by vaccination, we stimulated FACS sorted CD4 and CD8 T cells with spike peptides for 24 hours and measured secreted factors using Luminex **(Figure 3H)**. CD4 T cells secreted elevated levels of cytokines (IL-6, IL-10), cytotoxic molecules (Granzyme A and Granzyme B), and costimulatory factor (sCD137; soluble 4-1BB) **(Figure 3I)**. Additionally, modest induction of IL-2 and IL-4 by CD4 T cells was measured **(Supp Figures 3E)**. No significant production of immune mediators was noted by CD8 T cells except for Perforin and modest levels of IFNγ **(Supp Figure 3F)**. Finally, enhanced secreted levels of apoptotic factor sFas was observed in both CD4 and CD8 T cells following vaccination **(Supp Figure 3G)**.

### Vaccination and infection induce different T cell clonal expansions

Next, we compared the changes in T cell clonal dynamics with infection or vaccination. Vaccination was associated with a shift towards increased CDR3 lengths **(Figure 4A)**. Infection was associated with expansion of large clones (>100 cells), while vaccination induced expansion of primarily small sized clones (2-3 cells) **(Figures 4B, Supp Figure 4A, B);** however these expansions were highly variable among different vaccinated individuals **(Supp Figure 4B)**. Finally, both vaccination and infection were associated with a drop in clonal diversity; however, this drop was more dramatic following infection **(Figure 4C)**.

**Figure 4:**
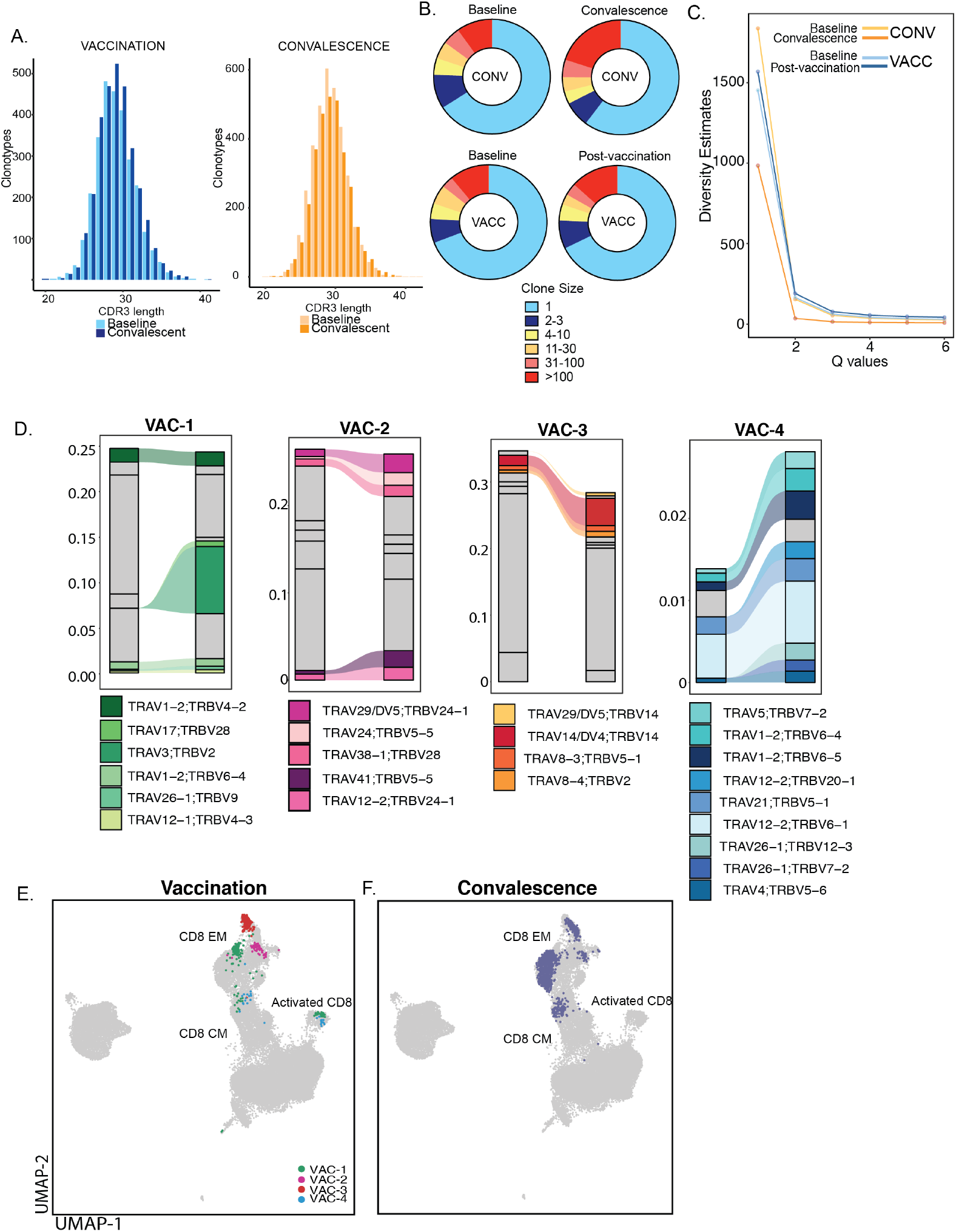
Clonal expansion of T cells following SARS-CoV-2 mRNA vaccination and infection. (A) CDR3 distribution of T cell clones following vaccination (left) and infection (right) (B) Pie charts representation of distribution of T cell clone sizes following vaccination and infection (C) Diversity profiles of T cells following vaccination and infection - y axis represents Hill diversity, interpreted as the effective number of clonotypes within the dataset. (D) Clonotype tracking in four volunteers two weeks following the second dose of mRNA vaccine. Only the top 10 clones post vaccination with evidence of clonal expansion following vaccination are highlighted. (E-F) UMAP projection of top 10 expanded clones following (E) vaccination (each volunteer highlighted) and (F) convalescence. Supp Figure 1: Experimental design and humoral response to vaccination and infection

We next compared biases in TCRα and TCRβ gene usage by comparing repertoire assignments post convalescence and vaccination with their respective baselines. Within TCRα, we observed limited overlap between vaccination and infection groups with a positive bias towards TRAV39 (p=0.0003 with convalescence; p=0.23 with vaccination), TRAV29/DV5 (p=0.3 with convalescence; p=0.09 with vaccination), TRAV21 (p=0.02 with convalescence; p=0.36 with vaccination) **(Supp Figure 4C)**, TRBV10 (p=0.1 with convalescence; p=0.0.03 with vaccination), and TRBV12 (p=0.09 with convalescence; p=0.2 with vaccination) **(Supp Figure 4D)**. However, convalescence and vaccination preferentially enriched distinct TCRs - TRAV29/DV5; TRBV5-1 and TRAV29/DV5; TRBV6-5 with convalescence **(Supp Figures 4E and 4F)** and TRAV29/DV5; TRBV11-2, TRAV29/DV5; TRBV7-9, and TRAV12-2; TRBV6-2 with vaccination **(Supp Figure 4G)**. We observed diverse patterns of clonal expansion with few clones that expanded dramatically in Vac 1-3 while several smaller clones with limited expansion were detected in Vac4 **(Figure 4D)**. Finally, the T cell clones that expanded with vaccination or convalescence occupied distinct space within the UMAP **(Figures 4E and 4F)**. Interestingly, the top expanded clones following vaccination were mostly CD8 EM with smaller involvement of CD8 CM and activated CD8 T cells **(Figures 1B and 4E).** However, expanded clones within convalescent individuals included both CD8 EM and CD8 CM **(Figures 1B and 4F)**.

## DISCUSSION

The establishment of immunity against severe acute respiratory syndrome coronavirus 2 (SARS-CoV-2) has become a central focus of current research efforts. Natural immunity following infection and vaccine-generated immunity provide two different pathways to immunity against the disease. mRNA vaccines have demonstrated significant protection against severe COVID-19 disease. Findings from human trials of Pfizer/BioNTech and Moderna suggest 95% maximal protection within 1-2 months after the second vaccine dose, including against several circulating variants of concern (9, 10). Recommendations from CDC indicate that individuals are not fully protected until 2 weeks after the second dose of vaccine (11). In this study, we investigated the cellular changes in circulating T and B cells induced by SARS-CoV-2 mRNA vaccination in four individuals before vaccination and 2 weeks after the second dose and compared their clonal adaptations in individuals who developed natural immunity against the virus.

The presence of neutralizing antibodies is currently used as a surrogate indicator of immunity. Both mRNA vaccines induce potent and durable neutralizing antibodies as early as 10 days (5) and last as long as seven months after the first dose of vaccination (12). Our data suggests that while neutralizing titers after the first dose are comparable to those observed in recovered individuals, levels of neutralizing titers are significantly higher than both groups following the second vaccination dose. However, both vaccinees and convalescent subjects shared several key B cell adaptations. For example, flow analysis revealed a reduction in naïve but expansion of memory B cells in both groups. Additionally, scRNA-seq analysis revealed a reduction in IgA+ (*IGHA1*) memory B cells following vaccination, as recently described in COVID-19 recovered individuals (13). Within PBMC from SARS-CoV-2 naive individuals, we observed a modest expansion of plasmablasts (in 3 out of 4 subjects), and a significantly elevated presence of Spike-reactive B cells (in all 4 subjects) following vaccination, suggesting establishment of durable memory and potential recall responses to infection.

Clonal analysis of memory B cells revealed expansion of small sized clones in recovered individuals, and to a lesser extent in vaccinated individuals. This is in line with recent reports of limited evidence of somatic hypermutation in spike-binding memory B cell clones in vaccinated individuals one week after second dose (5). We argue that this is likely because our sampling captured very early events in B cell responses and assessment at later time points would reveal more clones that emerge from prolonged B cell evolution within the germinal center as observed in recovered individuals (14, 15). Supporting this hypothesis, a recent study analyzing lymph node aspirates of vaccinated individuals observed S-binding GC B cells and plasmablasts for at least 15 weeks after first dose, further expanding after second dose (16). Interestingly, circulating IgG and IgA secreting spike specific plasmablasts peaked one week after the second dose and then declined, becoming undetectable three weeks later (16). This is in line with our findings, where we observed increased, but weak enrichment of CD27+CD38++ plasmablasts in blood 2 weeks following second dose. Finally, mRNA vaccine induced GC B cells are at near peak frequencies for at least 12 weeks after second dose (16), suggesting future assessment of changes in B cell repertoire at least 2-3 months following vaccination.

The characteristics of cellular immunity induced by mRNA vaccines still remain unclear. Early studies testing mRNA vaccine efficacy have demonstrated robust antigen specific Th1 responses following two doses of vaccines (7). Our single cell analyses suggest an expansion of activated CD4 T cells (CD38+HLA-DR+) but not CD8 T cells in every vaccinee two weeks after the second dose. In the presence of lower neutralizing antibodies following the first vaccine dose, rapid induction of SARS-CoV-2 specific CD4 T cells in SARS-CoV-2 naive individuals has been argued to play a role in protection after the first dose of vaccine (17). This is in contrast to antigen specific CD8 T cells which develop gradually and reach maximal levels only after the second dose (17). Indeed, overnight stimulation of PBMC with overlapping 15-mers covering full length spike protein revealed enrichment of polyfunctional CD4 T cells in all individuals (4 of 4) following the second dose. Cytokine analysis of stimulated CD4 T cells also suggested a robust effector response to spike peptides. In contrast, cytokine producing CD8+ T cells were observed in only 2 of the 4 individuals, and effector responses to spike peptides were weak, suggesting a delay in development of effector CD8 T cell response. Our findings are in line with early efficacy studies and recent follow up studies both one-week post booster and up to seven months post dose 1, where the magnitude of CD8 T cell responses (measured using both AIM assay and intracellular cytokine responses) was both variable and several fold lower in comparison to CD4 responses (7, 17). To some extent this is not surprising, given that the most immunodominant epitopes recognized by CD8 T cells in COVID-19 patients are contained in ORF1 and not spike protein (18). Nevertheless, whether the strength of early CD4 T cell response and/or variability in CD8 T cell responses in vaccinees is predictive of durable neutralizing titers and/or long term memory B cell responses is yet to be evaluated. Interestingly, persistence of antigen specific CD4 and CD8 T cells 7 months after the first dose (12) supports the hypothesis that mRNA vaccine induces durable CD4+ and CD8+ T cell responses capable of contributing to future recall responses.

Both vaccination and convalescence enriched T cell clones with longer CDR3s, though this shift was more prominent with vaccination. However, infection induced a sharper drop in diversity of T cell repertoire compared to vaccination. Surprisingly, clonal tracking analysis revealed expansion of effector memory CD8 T cells in all subjects, albeit the magnitude of expansion being very weak in one aged individual included in this study. This is in contrast to very limited expansion within CD4 T cells following the second dose of vaccination. We posit that analysis of T cell clones as early as 2 weeks post dose 1 would allow for better assessment of clonal expansion within CD4 T cells, which we might have missed given our sampling window. Alternatively, future studies will have to evaluate CD4 T cell clones at various time points following *in vitro* enrichment of antigen specific clones. Expansion of effector memory CD8 T cell clones following vaccination, however, is in line with what has been reported with natural infection (13, 19, 20). Despite the significant expansion of CD8 T cell clone, frequency of S-specific CD8 T cells was small and variable. This discrepancy could be due to bystander activation. Expanded T cell clones with vaccination and infection occupied distinct space on single cell maps, highlighting differences in the breadth of the epitopes recognized in vaccinated compared to infected individuals (18).

Interestingly, early post boost induction of spike specific memory B cells has been shown to correlate negatively with age following mRNA vaccination (5). Incidentally, the subject with the lowest CD8 T cell expansion and RBD binding antibodies in this study is also the oldest subject in our cohort (VAC-4). This is in line with data from clinical trials showing lower neutralizing responses after 100 ug dose and faster waning of the response following low dose (25 ug) of mRNA-1273 (12, 21) in the elderly. Furthermore, natural infection has been shown to impair SARS-CoV-2 specific priming of CD8 T cells in the elderly (22, 23). Whether that defect extends to vaccine induced early CD4 T cell responses or subsequent CD8 T cell expansion remains to be seen. Our study, however, was limited by sample size to draw definitive conclusions on weakening of vaccine responses in the aged. Moreover, it is still unclear what magnitude of neutralizing response confers protective immunity to the virus.

Limitations of our study include small sample size, and restriction to participants receiving mRNA vaccine. Due to limited blood samples collected, we were unable to perform additional analysis such as phenotyping of circulating T follicular helper cells (cTfh) following vaccination. Given such few memory B cells from each subject, we were unable to perform rigorous somatic hypermutation analysis at the single cell level. Finally, future studies will have to focus on long-term protection (both cellular and humoral) of two doses of mRNA vaccine against the numerous variants of SARS-CoV-2 and mechanisms of decline in quality of protection (if any) in the elderly.

## METHODS

### Ethics Statement

This study was approved by the University of California Irvine Institutional Review Boards. Informed consent was obtained from all enrolled subjects.

### Study Participants and Experimental Design

All participants in this study were healthy and did not report any comorbidities. All vaccines (VACC group) received either the Pfizer (BNT162b2) or the Moderna (mRNA-1273) mRNA-based vaccines. Blood was collected at 3 time points: pre-vaccine baseline, two weeks post-primary vaccine (dose 1), and two weeks post prime-boost vaccination (dose 2). Blood collected at two-weeks post primary vaccine (dose 1) was used only for serological experiments. Baseline and post-vaccination samples were analyzed for cellular and humoral response to the vaccine. For convalescent subjects (CONV group), blood samples collected pre-exposure to SARS-CoV-2 (Baseline) and ~30 days post convalescence were included in the analysis. These subjects experienced asymptomatic/mild COVID-19. Detailed characteristics of participants and experimental breakdown by sample is provided in Supp. Table 1.

### Plasma and Peripheral Blood Mononuclear Cells (PBMC) Isolation

Whole blood samples were collected in EDTA vacutainer tubes. PBMC and plasma samples were isolated after whole blood centrifugation 1200 g for 10 minutes at room temperature in SepMate tubes (STEMCELL Technologies). Plasma was stored at −80°C until analysis. PBMC were cryo-preserved using 10% DMSO/FBS and Mr. Frosty Freezing containers (Thermo Fisher Scientific) at −80C then transferred to a cryogenic unit 24 hours later until analysis.

### Measuring antibody responses

RBD end-point titers were determined using standard ELISA and plates were coated with 500 ng/mL SARS-CoV-2 Spike-protein Receptor-Binding Domain (RBD) (GenScript, Piscataway NJ) Heat inactivated plasma (1:50 in blocking buffer) was added in 3-fold dilutions. Responses were visualized by adding HRP-anti-human IgG (BD Pharmingen) followed by the additional of Phenylenediamine dihydrochloride (ThermoFisher Scientific). ODs were read at 490 nm on a Victor3 ™ Multilabel plate reader (Perkin Elmer). Batch differences were minimized by normalizing to positive control samples run on each plate.

Focus Reduction Neutralization Titer (FRNT) was measured using heat-inactivated plasma serially diluted (1:3) in HyClone Dulbecco’s Modified Eagle’s Medium (DMEM) supplemented with 10mM of HEPES buffer. The diluted plasma was pre-incubated with SARS-CoV-2 (100 PFU) for 1 hour before being transferred onto Vero E6 cells (ATCC C1008) seeded in a 96-well plate followed by overlay using 1% methylcellulose (Sigma Aldrich). After 24 hours, the medium was carefully removed, and the plates were fixed. Number of infected foci was determined using anti-SARS-CoV-2 Nucleocapsid antibody (Novus Biologicals NB100-56576) and HRP anti-rabbit IgG antibody (BioLegend). Plates were developed using True Blue HRP substrate and imaged on an ELISPOT reader. Each plate included a positive and a negative control. The half maximum inhibitory concentration (IC50) was calculated by non-linear regression analysis using normalized counted foci on Prism 7 (Graphpad Software). 100% of infectivity was obtained normalizing the number of foci counted in the wells derived from the cells infected with SARS-CoV-2 virus in the absence of plasma.

### Adaptive immune phenotyping

Frozen PBMCs were thawed, washed in FACS buffer (2% FBS, 1mM EDTA in PBS) and counted on TC20 (Biorad) before surface staining using the following panel: CD4 (OKT4, BioLegend), CD8b (2ST8.5H7, Beckman Coulter), CD28 (CD28.2, eBioscience), CD95 (DX2, eBioscience), CD20 (2H7, eBioscience), IgD (IA6-2, BioLegend), CD27 (M-T271, BioLegend), and CD38 (AT1, Stemcell Technologies). Dead cells were excluded using the Ghost Dye Red 710 (Tonbo). T cell phenotyping was conducted using an additional panel of antibodies – CD4 (OKT4, Biolegend), CD8b (2ST8.5H7, Beckman Coulter), CCR7 (G043H7, Biolegend), CD45RA (T6D11, Miltenyi Biotec), CD38 (HIT2, Tonbo Biosciences), CD27 (O323, Biolegend), HLA-DR (L243, Biolegend), CD69 (FN50, Biolegend), and PD-1 (EH12.2H7, Biolegend). All samples were acquired on the Attune NxT acoustic focusing cytometer (Life Technologies). Data were analyzed using FlowJo v10 (TreeStar, Ashland, OR, USA).

### Antigen specific T cell responses

Approximately 1X10e6 PBMC were stimulated with 1 ug of each of the SARS-CoV-2 peptide pool 5 (S protein) or anti CD3 (positive control) in 96 well plates for 24 at 37 C and 5% CO_2_. Plates were spun, surface stained using an antibody cocktail containing CD4 (OKT4, BioLegend), CD8b (2ST8.5H7, Beckman Coulter), CD28 (CD28.2, eBioscience), CD95 (DX2, eBioscience). Cells were washed, fixed and stained intracellularly using TNFɑ (MAB11, eBioscience), IL-2 (MQ1-17H12, Biolegend), and IFNγ (4S.B3, eBioscience). Samples were analyzed on Attune NxT Flow cytometer (ThermoFisher). Data were analyzed on FlowJo (BD Biosciences).

Approximately 5×10e4 CD4 and CD8 T cells were sorted and stimulated with 1 ug of the SARS-CoV-2 peptide pools Pool 5 (S protein) or anti CD3 (positive control) for 16h at 37C and 5% CO_2_. Plates were spun, and supernatants collected and stored in −80C. Immune mediators in supernatants were measured using a Milliplex MAP Human CD8+ T cell 17-plex magnetic bead panel measuring GM-CSF, sCD137, IFNγ, IL-10, Granzyme A, Granzyme B, IL-13, sFas, IL-2, IL-4, IL-5, sFasL, MIP-1ɑ, MIP-1β, TNFɑ, and Perforin per manufacturer’s instructions and run on Magpix (Luminex Corp, Austin TX). Standard curves were fit using 5P-logistic regression on XPonent software (Luminex Corp, Austin TX)

### Antigen Specific B cells

To detect antigen specific B cells, ~ 5×10e5 PBMC were stained with 100 ng of full length biotinylated spike protein (Sino Biological) pre-incubated with Streptavidin-BV510 (Biolegend) at 2:1 ratio for 1 h at 4C to ensure maximum staining quality before surface staining with CD20-FITC (2H7, Biolegend) for an additional 30 minutes. Streptavidin PE (Biolegend) was used as a decoy probe to gate out SARS-CoV-2 nonspecific streptavidin binding. Samples were washed twice and resuspended in 200 uL FACS buffer before being analyzed on Attune NxT (Life Technologies).

### FACS for repertoire analysis

Cryopreserved PBMC from each person (n=4 for pre- and post-vaccine samples; n=3 for baseline and convalescent samples) were thawed, washed, and stained with 1 ug/test cell-hashing antibody (TotalSeq C0251, C0254, C0256, C0260, clones LNH-95, 2M2, BioLegend) for 30 minutes at 4C. Samples were washed three times in ice cold PBS supplemented with 2% FBS and sorted on the FACSAria Fusion (BD Biosciences) with Ghost Dye Red 710 (Tonbo Biosciences) for dead cell exclusion, and then CD4, CD8, CD28, CD95, CD38, CD27, and IgD to sort memory CD4, CD8 T cells, memory B cells, and plasmablasts. Live, sorted cell populations were counted in triplicates on a TC20 Automated Cell Counter (BioRad) and pooled into 4 samples (pre-vaccine, post-vaccine, baseline, and convalescent).

### 5’ single cell RNA sequencing

Pooled cells were resuspended in ice cold PBS with 0.04% BSA in a final concentration of 1800 cells/uL. Single cell suspensions were then immediately loaded on the 10X Genomics Chromium Controller with a loading target of 26,000 cells. Libraries were generated using the Chromium Next Gem Single Cell 5’ Reagent Kit v2 (Dual Index) per manufacturer’s instructions with additional steps for the amplification of HTO barcodes and V(D)J libraries (10X Genomics, Pleasanton CA). Libraries were sequenced on Illumina NovaSeq with a sequencing target of 30,000 reads per cell RNA library, 5,000 reads per cell HTO barcode library, and 5,000 reads per cell for V(D)J libraries.

### Single cell RNA-Seq data analysis

Raw reads were aligned and quantified using the Cell Ranger Single-Cell Software Suite with Feature Barcode addition (version 4.0, 10X Genomics) against the GRCh38 human reference genome using the STAR aligner. Downstream processing of aligned reads was performed using Seurat (version 4.0) (38). Droplets with ambient RNA (cells fewer than 200 detected genes), dying cells (cells with more than 20% total mitochondrial gene expression), and cells expressing both a TCR and BCR clonotype were excluded during initial QC. Data normalization and variance stabilization was performed on the integrated object using the *NormalizeData* and *ScaleData* functions where a regularized negative binomial regression corrected for differential effects of mitochondrial gene expression levels. The *HTODemux* function was then used to demultiplex donors and further to identify doublets, which were then removed from the analysis. Dimension reduction was performed using *RunPCA* function to obtain the first 30 principal components followed by integration using Harmony. Clusters were visualized using the UMAP algorithm as implemented by Seurat’s *RunUMAP* function. Cell types were assigned to individual clusters using *FindMarkers* function with a fold change cutoff of at least 0.4. List of cluster specific markers identified from this study are cataloged in Supp Table 2. Differential expression analysis was performed using MAST using default settings in Seurat. All disease comparisons were performed relative to healthy donors from corresponding age groups. Only statistically significant genes (Log_10_(fold change) cutoff ≥ 0.25; adjusted p-value ≤ 0.05) were included in downstream analysis

### TCR and BCR analysis

TCR and BCR reads were aligned to VDJ-GRCh38 ensembl reference using Cell Ranger 4.0 (10X Genomics) generating sequences and annotations such as gene usage, clonotype frequency, and cell specific barcode information.

As an additional QC, only cells with one productive alpha and one productive beta chain were retained for downstream analyses. CDR3 sequences were required to have length between 5 and 27 amino acids, start with a C, and not contain a stop codon. Cells with both TCR and BCR (<0.1%) assignments were excluded from the analysis and all downstream analysis performed using the R package immunarch. Data were first parsed through *repLoad* function in immunarch, and clonality examined using *repExplore* function. Family and allele level distributions of TRA and TRB genes were computed using *geneUsage* function. Diversity estimates (Hill numbers) were calculated using *repDiversity* function and tracking of abundant clonotypes was performed using *trackClonotype* function.

Clonal assignments based on heavy and light chains were determined using change-o package in Immcantation portal. Briefly, heavy chain file was clonally clustered separately correct clonal groups assigned based on light chain data, removing cells associated with more than one heavy chain. Germline sequences were reconstructed using IgBlast. Gene usage, isotype abundance, and clonotype abundance were calculated using Alakazam package in Immcantation portal.

### Statistical analysis

Data sets were first tested for normality. All pairwise comparisons for readouts before/after vaccine and infection were tested using paired t-test. For comparisons involving multiple groups, differences were tested using one-way ANOVA followed by Holm Sidak’s multiple comparisons tests. P-values less than or equal to 0.05 were considered statistically significant. Values between 0.05 and 0.1 are reported as trending patterns.

## Author Contributions

S.S., S.A.L., and I.M. conceived and designed the experiments. S.S., S.A.L., B.D., and A.J. performed the experiments. S.S., S.A.L., and B.D. analyzed the data. S.S., S.A.L., and I.M. wrote the paper. All authors have read and approved the final draft of the manuscript.

## Acknowledgements

We are grateful to all participants in this study. We thank Dr. Jennifer Atwood for assistance with sorting and imaging flow cytometry in the flow cytometry core at the Institute for Immunology, UCI. We thank Dr. Melanie Oakes from UCI Genomics and High-Throughput Facility for assistance with 10X library preparation and sequencing. Aspects of experimental design figures were generated using graphics from Biorender.com.

## Data availability

The datasets supporting the conclusions of this article are available on NCBI’s Sequence Read Archive (SRA# pending).

---

**Supp Figure 1:**
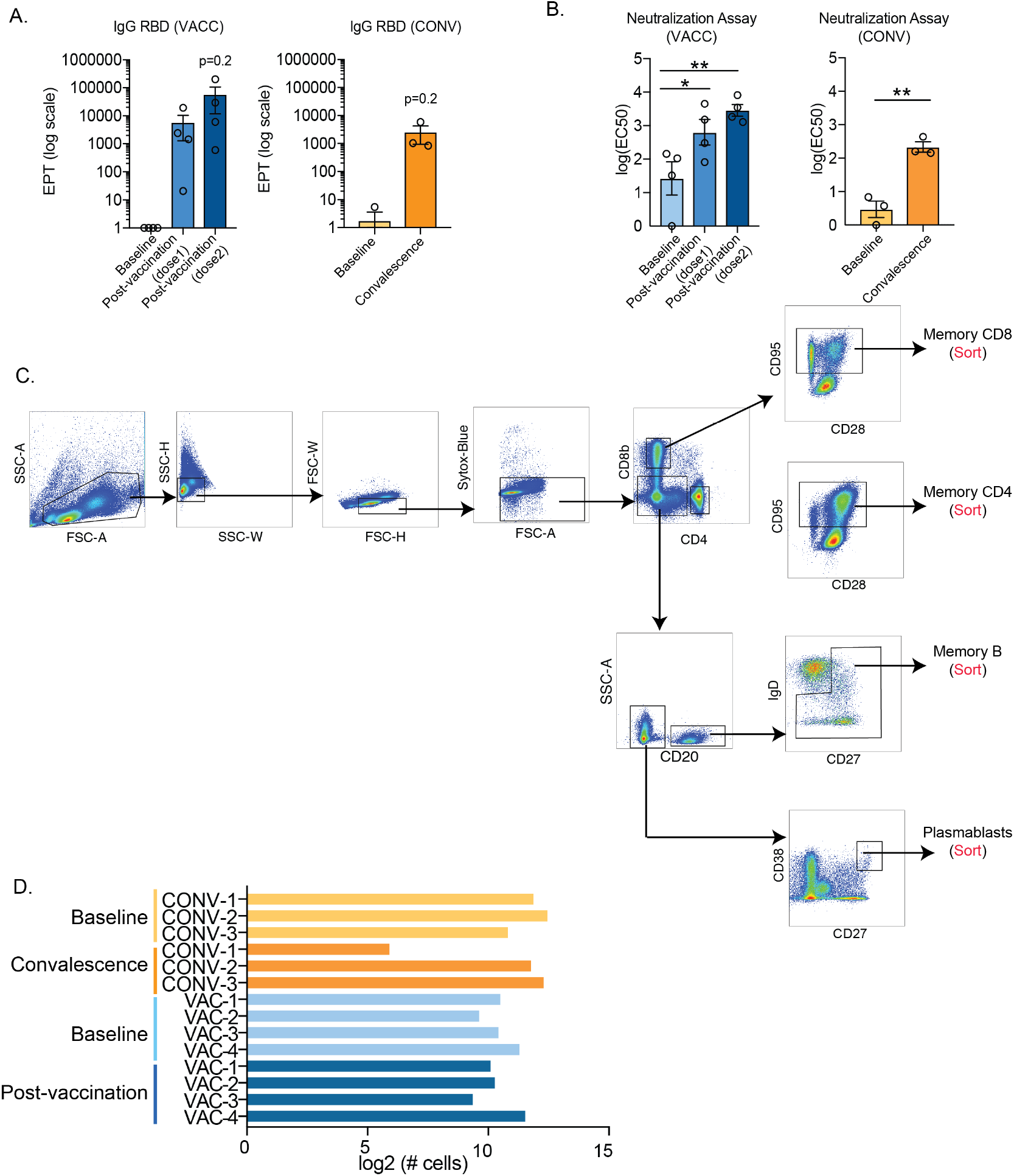
Experimental design and humoral response to vaccination and infection. (A) Bar plots comparing IgG Endpoint titers before and after vaccination (two weeks post dose 1 and dose 2) and before and after asymptomatic infection. (B) Bar plots comparing antibody neutralization of SARS-CoV-2 virus (IC50) before and after vaccination (two weeks post dose 1 and dose 2) and before and after asymptomatic infection. (C) Gating strategy for enrichment of memory T and B cells for single cell RNA and repertoire analysis. Cells were surface hashed using Total-Seq A antibodies and sorted based on surface CCR7 and CD45RA (for memory T cells), CD27 and IgD (for memory B cells), and CD27 and CD38 (for plasmablasts). (D) Sample wise breakdown of numbers of single cells recovered from each subject and used for downstream dimension reduction and clustering. Two-way comparisons were tested using unpaired t-test with Welch’s correction for group comparisons, [p-values: *− p<0.05; **−p<0.01]

**Supp Figure 2:**
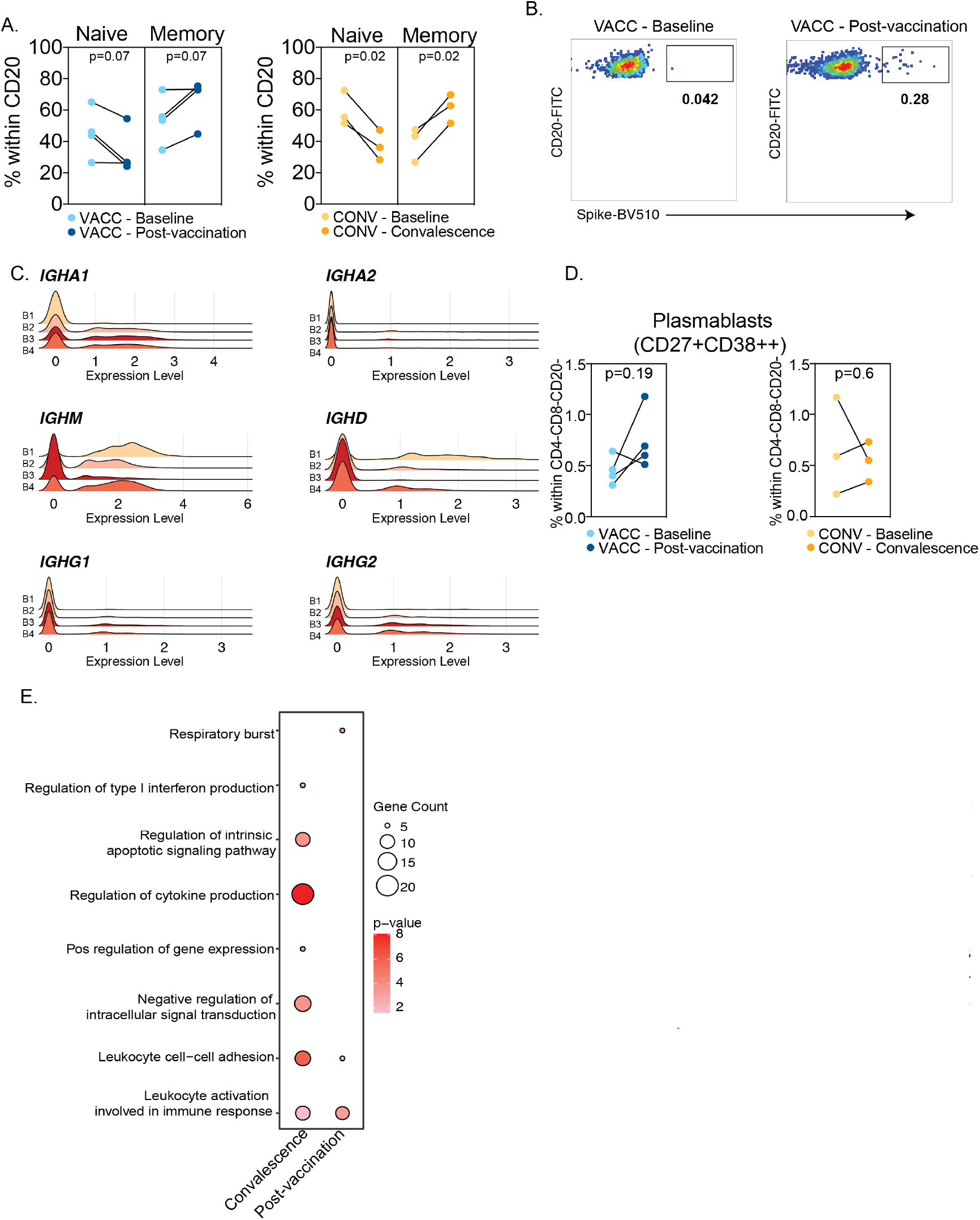
B cell adaptations with SARS-CoV-2 mRNA vaccine. (A) Frequencies of naive and memory B cells before and after vaccination (left) vs. before and after asymptomatic infection (right). (B) Representative plots for identification of Spike+ events within gated B cells. (C) Ridge plots of immunoglobulin genes across four memory B cell clusters identified using single cell RNA sequencing. (D) Frequencies of plasmablasts before and after vaccination (left) vs. before and after asymptomatic infection (right). (E) Bubble plots representing two-way functional enrichment of genes up-regulated with infection or vaccination in a direct comparison of memory B cells post-vaccination vs. convalescence. Size of the bubble represents the number of genes mapping to the term whereas color represents level of statistical significance. Two-way differences were tested using either ratio paired for matched comparisons.

**Supp Figure 3:**
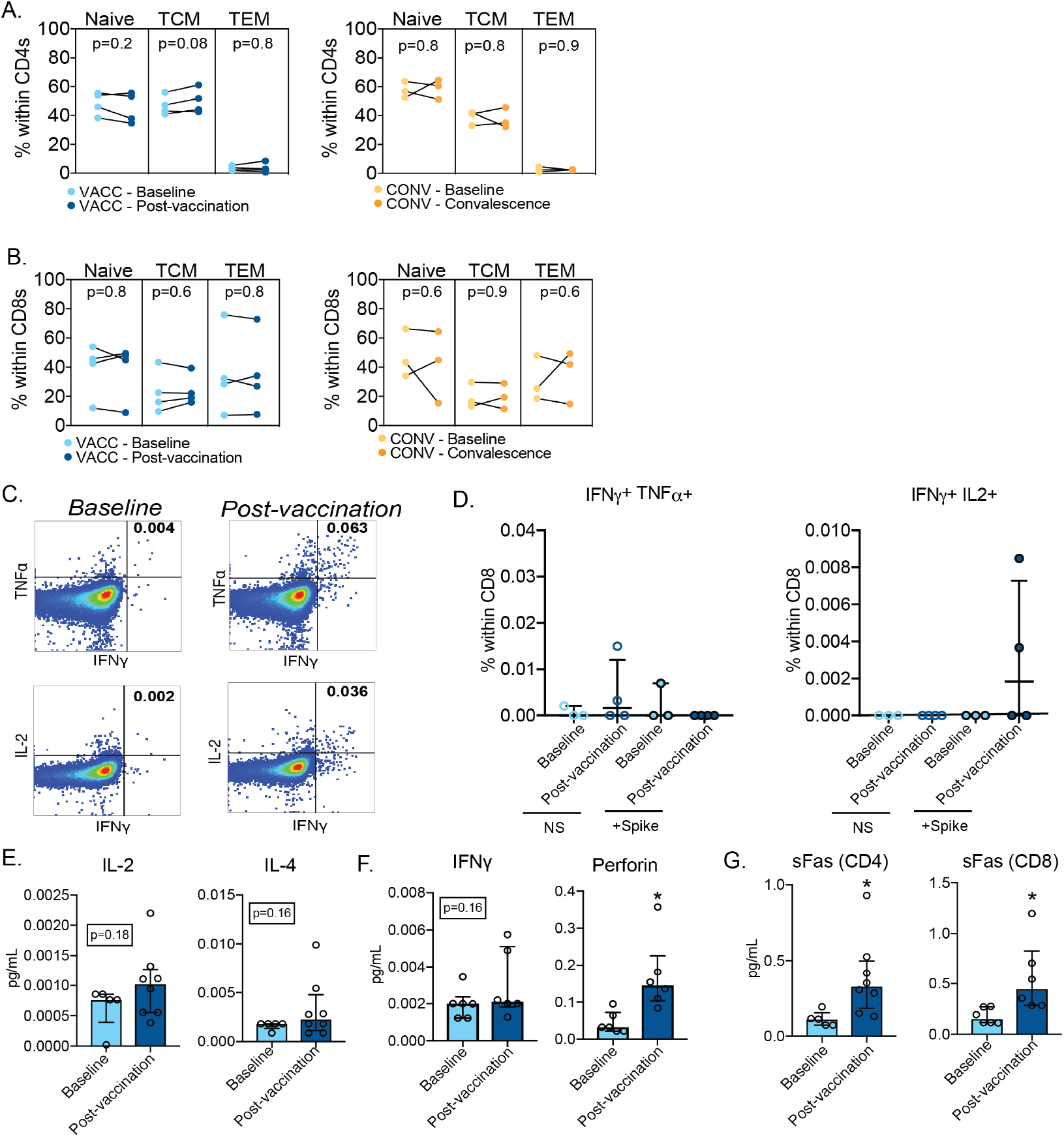
T cell adaptations with SARS-CoV-2 mRNA vaccination. (A-B) Frequencies of (A) CD4 and (B) CD8 T cells and their subsets before and after vaccination. (C) Gating strategy for identification of cytokine producing cells following in vitro stimulation of PBMC with SARS-CoV-2 overlapping spike peptides. (D) Polyfunctional CD8 T cell responses following overnight SARS-CoV-2 spike peptide stimulation measured using intracellular cytokine staining and flow cytometry. (E) Secreted levels of Th1 and Th2 cytokines by CD4 T cells following mRNA vaccination. (F) Secreted levels of IFNy and Perforin by CD8 T cells and (FI) sFAS molecule by CD4 and CD8 T cells following peptide stimulation before and after mRNA vaccine. Two-way comparisons were tested using either ratio-paired for matched comparisons or unpaired t-test with Welch’s correction for group comparisons. Four way comparisons were tested using one-way ANOVA followed by Flolm Sidak’s multiple hypothesis correction [p-values: *− p<0.05]

**Supp Figure 4:**
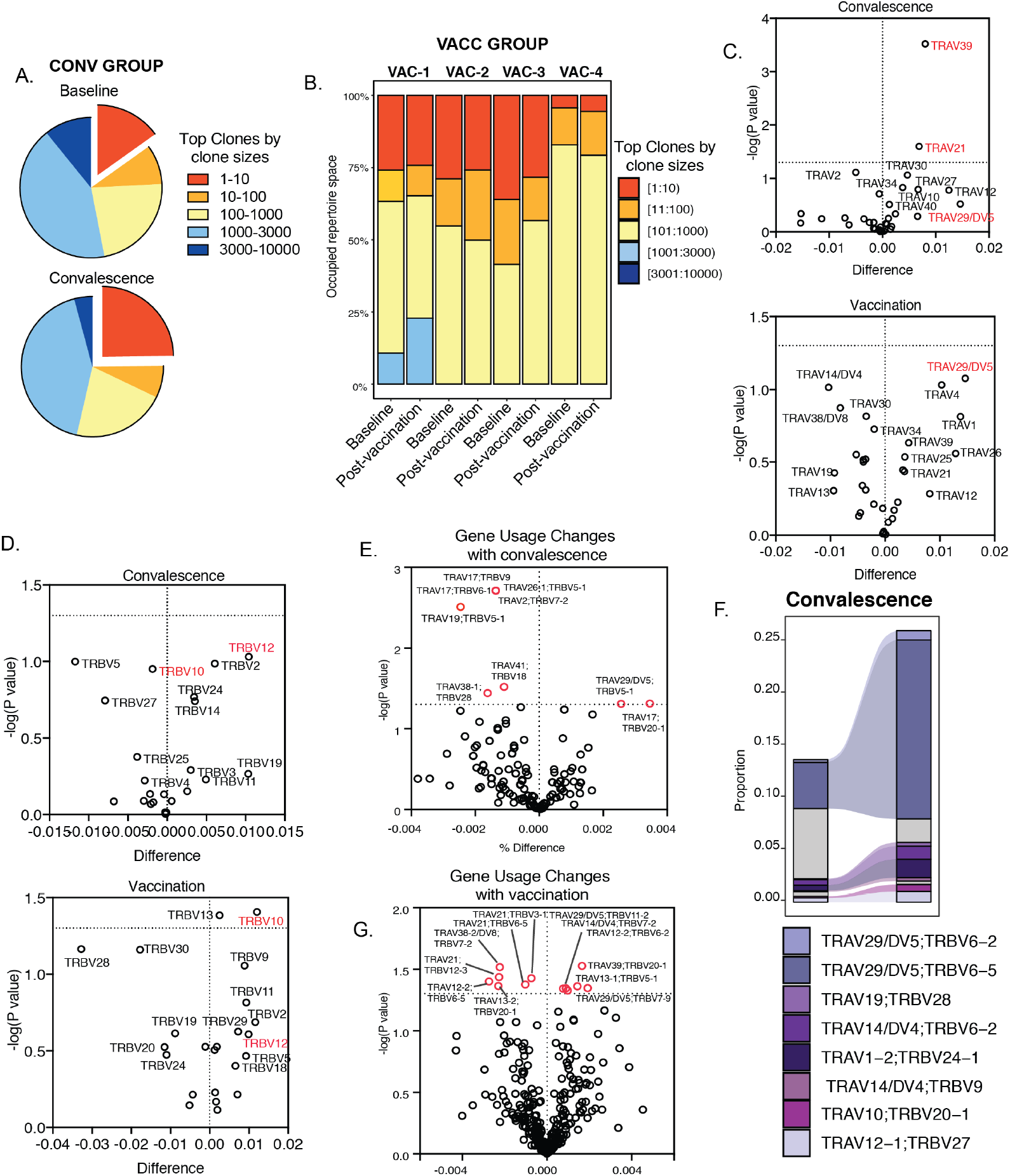
T cell clonal expansion with SARS-CoV-2 mRNA vaccine. (A) Pie charts demonstrating aggregate clonal space occupied by top clones (by size) before and after convalescence. (B) Stacked bars showing distribution of top clones before and after vaccination in four volunteers. (C-D) Volcano plots depicting (C) TCRa and (D) TCRp gene usage following convalescence (top panel, relative to pre-infection baseline) or vaccination (bottom panel, relative to pre vaccination baseline) at the gene family level. (E-F) Volcano plots depicting gene usage changes (TCRap combinations) following (E) convalescence (top panel, relative to pre-infection baseline) or (F) vaccination (bottom panel, relative to pre vaccination baseline). X axis represents the change in gene usage and Y axis represents p-value (−log 10). Statistical differences were tested using multiple t-tests. (G) Expanded T cell clones following convalescence. Only the top 10 clones post SARS-CoV-2 exposure with evidence of clonal expansion following infection are highlighted.

